# GIL: A Python package for designing custom indexing primers

**DOI:** 10.1101/2022.11.04.514296

**Authors:** Nicholas Mateyko, Omar Tariq, Xinyi E. Chen, Will Cheney, Asfar Lathif Salaudeen, Ishika Luthra, Najmeh Nikpour, Abdul Muntakim Rafi, Hadis Kamali Deghan, Cassandra Jensen, Carl de Boer

**Affiliations:** School of Biomedical Engineering of University of British Columbia, 2222 Health Sciences Mall, Vancouver, BC, V6T 1Z3

## Abstract

Next-generation sequencing allows samples to be multiplexed by adding a unique DNA index to each sample. Multiplexing greatly reduces the price of sequencing large numbers of samples, yet the minimum cost per sample remains high when using commercially available indexing kits and designing custom indexes is challenging. To address these issues, we created GIL (Generate Indexes for Libraries), a software tool that designs indexing primers for producing multiplexed sequencing libraries. GIL can be customized in numerous ways to meet user specifications, including index length, sequencing modality, color balancing, and compatibility with existing primers, and produces ordering and demultiplexing-ready outputs. GIL is written in Python and is freely available on GitHub at https://github.com/de-Boer-Lab/GIL. It can also be accessed as a web-application implemented in Streamlit at https://dbl-gil.streamlitapp.com.

## Introduction

Next-Generation Sequencing (NGS) has become a cornerstone of biology as a crucial method for data collection. The cost of sequencing has decreased with the refinement of sequencing technologies such that the cost of sequencing the whole human genome has decreased by a factor of nearly 3 million since 2003 (Check Hayden 2014). While advances in NGS technologies have facilitated remarkably low-cost acquisition of massive sequencing data, the minimum cost of NGS remains quite high (Schwarze et al. 2020). Multiplexing samples into a single-pooled library to reduce costs is a solution that has been explored since the earliest days of DNA sequencing (Church and Kieffer-Higgins 1988; Chee 1991). Multiplexing allows many samples to be sequenced in a single run, enabling researchers to take advantage of NGS’s low costs even when the desired sequencing depth for a single sample falls well below the scale of an NGS run (Smith et al. 2010). Multiplexing is accomplished by appending unique barcodes, or indexes, to each sample. The incorporation of these indexes by appropriately modified amplification primers has become commonplace over the past decade (O’Donnell et al. 2016). Compatible primers can be purchased as part of commercially distributed sample preparation kits, but at a considerable cost when large numbers of samples are prepared. Designing and ordering primers independently can reduce costs, but remains challenging due to the high cost associated with testing an indexing primer set and the many considerations one could account for when designing primers. A set of compatible indexing primers must be sufficiently dissimilar from one another for demultiplexing, avoid self-priming interactions, and be appropriately color balanced for the sequencing modality (Illumina 2018). Custom indexes would facilitate efficient large-scale sequencing at low costs.

Here we present GIL (Generate Indexes for Libraries), a user-friendly Python package that produces customizable sequencing primers in a ready-to-order and ready-to-demultiplex format. Users can provide custom adapter sequences, enabling index generation for any sequencing system or modality, and can create indexes of any length (e.g. greater than the standard 8 nt). Users can customize filtering to eliminate indexes that may cause issues in their setup (e.g. have repetitive sequences, or match existing primer sets). The generated order sheets can then be used to purchase primers at a fraction of the cost per sample of commercial library preparation kits.

## Methods

GIL is written in Python 3, and can be run both from the command line, or from an online Streamlit application. We designed GIL to create primers that add indexes to libraries by PCR. Input libraries must share a common adapter sequence on the ends, which can be added in multiple ways, including PCR for targeted sequencing, adapter ligation, or tagmentation, similar to the NEBNext® Multiplex Oligos for Illumina® and iTru approaches (Glenn et al. 2019).

**Figure 1.**
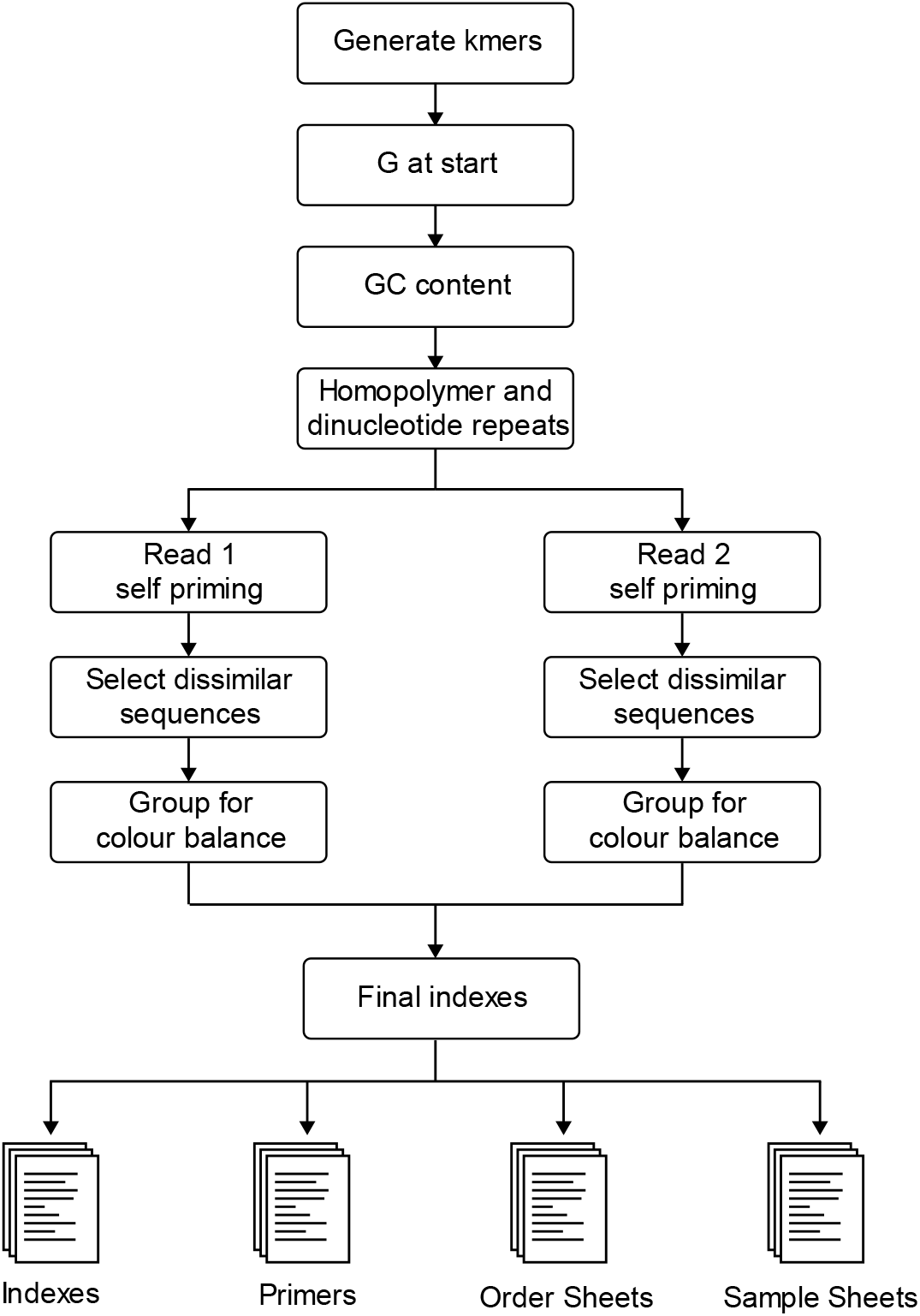
Overview of GIL index generation pipeline.

### Implementation

We first generate a set of indexes, either all k-mers for index length k ≤ 9, or a random sample of 5000 kmers for k > 9. We filter out indexes with undesirable qualities, find sequences from those that remain that are all sufficiently dissimilar to each other, and create a 96-well plate layout where the order of the barcodes on the plate maintains colour balance. The default filtering steps are as follows, although the parameters and filtering steps can all be customized:

1. **Remove indexes that start with G**. In Illumina sequencers that use 2-channel chemistry, G is not labelled with a fluorescent dye. Illumina warns that if the first two bases in the index are both G, “intensity is not generated” (Illumina 2018), and so indexes where the index read starts with G are removed.
2. **Remove indexes with extreme GC content**. We removed indexes with GC content ≤ 25% or ≥ 75%.
3. **Remove homopolymer and dinucleotide repeats**. We removed indexes with >2 homopolymer or dinucleotide repeats since simple repeats are associated with DNA synthesis errors.
4. **Remove indexes too similar to an existing index set (optional)**. In order for two sets of indexes to be compatible, they must be sufficiently distinct to uniquely identify the samples when demultiplexing. Users can provide an existing set of indexing primers (e.g. those that they use already or are commonly used at their sequencing facility), and all the indexes that are within 3 Levenshtein distance of the existing indexes are removed.
5. **Remove self-priming sequences**. We compute the Hamming distance between the reverse complement of the 8 bases on the 3’ end of the primer and all 8 nt windows that include the index sequence. If any of these distances are less than 3, the index is filtered out.
6. **Color balancing**. When only a few indexes are used in a sequencing run, color balance must be considered (Illumina 2018). We place indexes within a 96-well plate such that groups of four indexes along the rows of the plate are color balanced. This allows for multiplexing with as few as four consecutive indexes without having to consider color balance issues. We then place the generated indexes into the primer sequence context, which, by default, are designed to work with Illumina TruSeq. We designed the primers to have a Tm of 65 °C with NEB Q5 polymerase.

GIL has two main outputs. For each plate of indexes, GIL generates 1) an order sheet (CSV) that contains the well, primer name, and primer sequence columns in an order-ready format, and 2) a demultiplexing sample sheet in the standard Illumina format. Because some sequencers read index 2 in the reverse complement direction, two sample sheets are generated for each index plate, one with the index 2 column reverse complemented. Since a sample can be uniquely identified by the combination of index 1 and index 2 sequences, a single 96 well plate of each of index 1 and 2 primers can be used to index 962 (9216) samples that are compatible for pooling and sequencing together. However, most will not require so many indexing primers and so by default we create demultiplexing sample sheets where both index 1 and index 2 are unique and redundant, and could be used individually to demultiplex samples, enabling the detection and exclusion of index hopping reads (Farouni et al. 2020; van der Valk et al. 2020).

### Ordering primers

GIL was run with default parameters and generated three 96-well plates of TruSeq index primers. The 96 i5 and 96 i7 primers from the first generated plate (Supplemental File 1, Supplemental File 2) were ordered as 100 nmol oligos with standard desalting from IDT, including a single phosphorothioate bond between the last two bases on the 3’ end of the primers to prevent 3’-to-5’ degradation by DNA polymerase (Skerra 1992).

## Conclusion

After generating 3 plates of mutually compatible primers with default settings through GIL, we ordered primers, indexed samples (n=44), and sequenced them on the Illumina MiSeq Nano platform. Sequencing BCL files were demultiplexed successfully using the generated sample sheet (Supplemental File 3) with bcl2fastq software. All samples were present. Roughly 0.2% of reads within an index were thrown out due to deletions compared to 0.02% for a PhiX control, likely resulting from us opting for minimal primer purification.

### Comparison to existing software

Several programs for designing and testing sequencing indexes exist. EDITTAG (Faircloth and Glenn 2012), BARCRAWL and BARTAB (Frank 2009), and DNABarcodes (Buschmann and Bystrykh 2013) are all freely available tools for producing custom indexes for multiplexed sequencing. Each of these solutions provide some of the functionality found in GIL, however GIL has several advantages over existing solutions. For instance, GIL allows the user to consider self-priming interactions with the constant flanking primer sequence, the presence of a 5’ G, and dinucleotide repeats. Furthermore, GIL can produce primers with arbitrary constant regions flanking the indexes, enabling atypical uses or non-Illumina platforms. Finally, GIL is much easier to use, providing primers in an order-ready format, the files needed for demultiplexing (bcl2fastq), and a graphical interface.

**Table 1.**
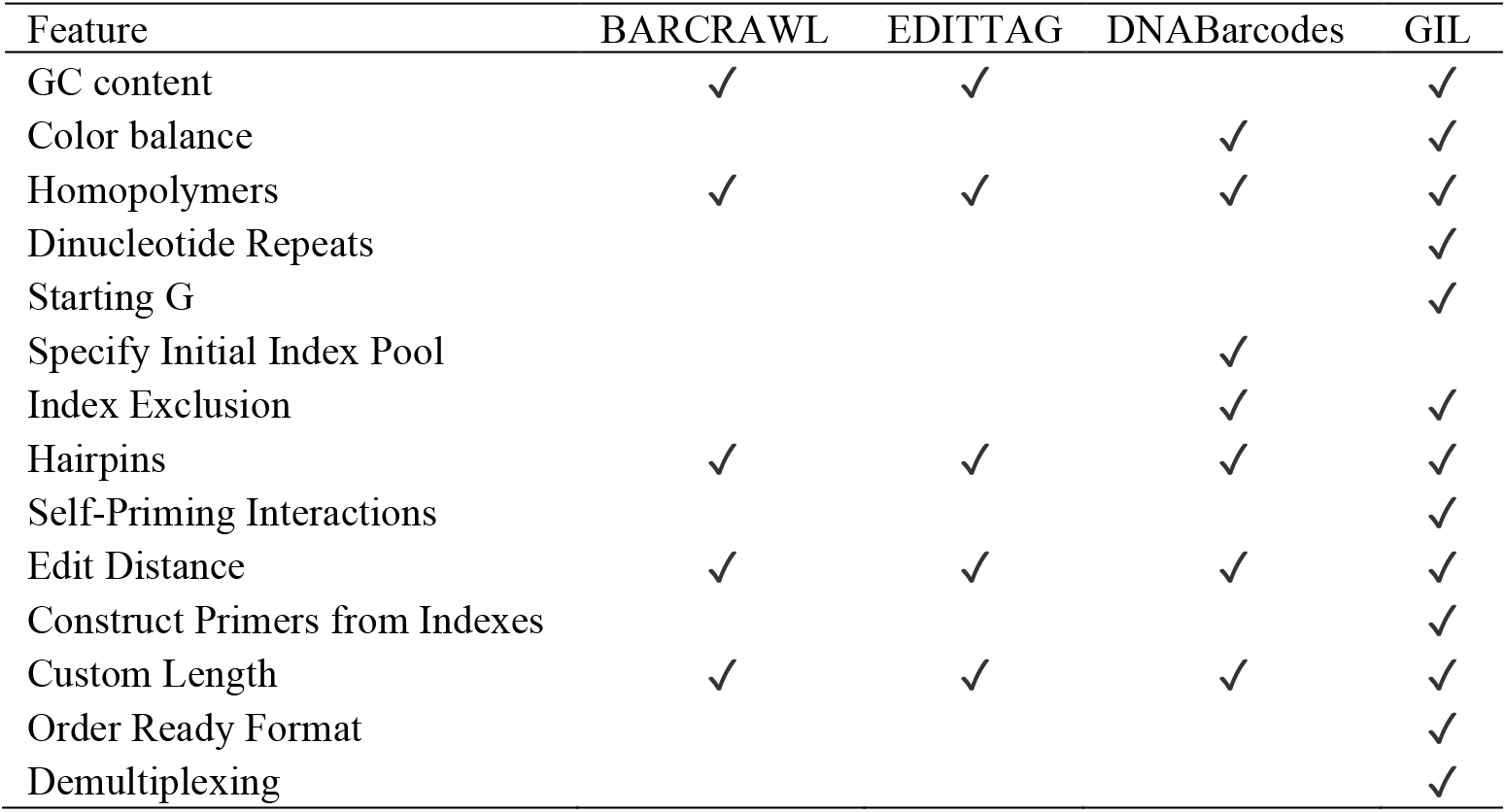
Comparison of GIL to existing software

| Feature | BARCRAWL | EDITTAG | DNABarcodes | GIL |
| --- | --- | --- | --- | --- |
| GC content | ✓ | ✓ |  | ✓ |
| Color balance |  |  | ✓ | ✓ |
| Homopolymers | ✓ | ✓ | ✓ | ✓ |
| Dinucleotide Repeats |  |  |  | ✓ |
| Starting G |  |  |  | ✓ |
| Specify Initial Index Pool |  |  | ✓ |  |
| Index Exclusion |  |  | ✓ | ✓ |
| Hairpins | ✓ | ✓ | ✓ | ✓ |
| Self-Priming Interactions |  |  |  | ✓ |
| Edit Distance | ✓ | ✓ | ✓ | ✓ |
| Construct Primers from Indexes |  |  |  | ✓ |
| Custom Length | ✓ | ✓ | ✓ | ✓ |
| Order Ready Format |  |  |  | ✓ |
| Demultiplexing |  |  |  | ✓ |

## Acknowledgements

We thank Marjan Barazandeh for their help in testing GIL’s implementation.

